# Cryopreservation of mosquito microbiota for use in microcosm experiments

**DOI:** 10.1101/2022.07.13.499917

**Authors:** Serena Y. Zhao, Grant L. Hughes, Kerri L. Coon

**Author notes:** Corresponding author: Kerri L. Coon.

## Abstract

Mosquitoes develop in a wide range of aquatic habitats containing highly diverse and variable bacterial communities that shape both larval and adult traits, including the capacity of adult females of some mosquito species to vector disease-causing organisms to humans. However, while most mosquito studies control for host genotype and environmental conditions, the impact of microbiota variation on phenotypic outcomes of mosquitoes is often unaccounted for. The inability to conduct reproducible intra- and inter-laboratory studies of mosquito-microbiota interactions has also greatly limited our ability to identify microbial targets for mosquito-borne disease control. Here, we developed an approach to isolate and cryopreserve microbial communities derived from mosquito larval rearing environments in the lab and field. We then validated the use of our approach to generate experimental microcosms colonized by standardized lab- and field-derived microbial communities. Our results overall reveal minimal effects of cryopreservation on the recovery of bacteria when directly compared with isolation from non-cryopreserved fresh material. Our results also reveal improved reproducibility of microbial communities in replicate microcosms generated using cryopreserved stocks over fresh material. Altogether, these results provide a critical next step toward the standardization of mosquito studies to include larval rearing environments colonized by defined microbial communities. They also lay the foundation for long-term studies of mosquito-microbe interactions and the identification and manipulation of taxa with potential to reduce mosquito vectorial capacity.

## 1. Introduction

The community of microbes (or ‘microbiota’) present in the aquatic environments where mosquito larvae develop can have profound impacts on mosquito biology by modulating larval growth and development and, consequently, adult survival, reproduction, and the competency of adult female mosquitoes to transmit human pathogens (reviewed in (1)). However, the enormous diversity and complexity in microbiota composition between different larval rearing environments in both the laboratory and field has to date made it difficult to assign functions to specific community members (1). The impact of microbiota variation on mosquito phenotypes important for vectorial capacity is also likely driven by interactions with mosquito genotype and environmental factors like diet and temperature (2–16), as is well-documented in vertebrate models (17–31). However, while experiments are often performed in a controlled and standardized environment using inbred mosquito strains, the microbiota is often not taken into account as a potential source of variation and likely underlies why mosquitoes may respond differently to an intervention in one study compared to another (32–54).

We recently developed an approach to successfully isolate and transfer complete microbial communities within and between adult mosquitoes of different donor and recipient species (55). We then demonstrated the utility of this approach to study the factors shaping microbiome acquisition and assembly in mosquitoes and the mechanisms by which specific microbial taxa and assemblages contribute to different mosquito traits under controlled conditions (55). However, while these results highlight the value of expanding tools to manipulate the microbiota in mosquitoes, important questions remain regarding (*i*) the utility of microbiome transplantation approaches to establish reproducible communities in mosquito larval rearing environments, which harbor microbial communities that are much more complex than those in adult mosquitoes (5,8,52,56–62), and (*ii*) how microbiota transplantation efficacy may be shaped by long-term preservation of donor microbial communities (*e.g*., via cryogenic freezing), which is absolutely necessary to facilitate long-term studies and intra- and interlaboratory comparisons but may introduce additional variability via impacts on bacterial viability and recovery (63–68).

In this study, we developed an approach to isolate and cryopreserve microbiota from mosquito larval rearing environments in the laboratory and field. We then validated the use of this approach to generate experimental microcosms colonized by standardized lab- and field-derived microbial communities. Our results overall reveal minimal effects of cryopreservation on the recovery of bacteria when directly compared with isolation from non-cryopreserved fresh material. These findings also reveal improved reproducibility of microbial communities in replicate microcosms generated using cryopreserved stocks over fresh material. Altogether, our work provides a critical next step toward the standardization of mosquito studies to include larval rearing environments colonized by defined microbial communities. They also lay the foundation for long-term studies of mosquito-microbe interactions and the identification and manipulation of taxa with potential to reduce mosquito vectorial capacity.

## 2. Methods

### (a) Laboratory colony, microbiota isolation and cryopreservation

Microbiota used in this study were derived from a laboratory colony of *Aedes aegypti* mosquitoes (Liverpool strain), which is conventionally reared at a constant temperature of 27°C, relative humidity (RH) of 70%, and photoperiod of 16-h light: 8-h dark (69). In brief, adults are housed in stainless-steel cages (BioQuip, Rancho Dominguez, CA, USA) and provided 10% sucrose in water *ad libitum*. Adult females are blood-fed once per week to promote egg laying using defibrinated sheep blood (Hemostat Laboratories, Dixon, CA, USA) via an artificial membrane feeder. Four days after blood feeding, eggs are collected from colony cages and stored at 27°C and 70% RH. Eggs are then hatched in enamel pans containing deionized water and larvae are fed a standard diet consisting of TetraMin Tropical Flakes Fish Food (Tetra, Melle, Germany) until pupation.

We isolated the microbiota present in the conventional larval rearing environment of our *A. aegypti* colony by sampling water from four replicate rearing pans containing larvae that had molted to the final (fourth) instar. For each pan, 500-ml of water was collected in sterile conical tubes (Thermo Fisher Scientific, Waltham, MA, USA) and used to generate the following resources for downstream experiments and sequencing as follows (Fig. 1): (*i*) 50-ml of water was immediately centrifuged at high speed (21 130 x *g*) for 20-min, supernatant removed, and pellet stored at −20°C for downstream DNA isolation and sequencing to characterize microbiota diversity, (*ii*) 200-ml of water was stored unprocessed at 4°C for use in downstream experiments, and (*iii*) the remaining volume (250-ml) was serially diluted, centrifuged at low speed (3220 x *g*) for 20-min, resuspended in 5-ml sterile PBS (1X) containing 20% glycerol, and frozen overnight at −80°C to produce a total of 24 cryopreserved stocks at variable cell densities (~10^8^-10^3^ bacterial cells per ml). Cell density was estimated by counts of colony-forming units (CFU) on R2A agar plates. In brief, aliquots of each cryopreserved microbiota stock were diluted to 10^-4^ and 50-ul of the diluted suspensions were used for plating in triplicate. Cell densities for each stock were then estimated using the average CFU count among replicate plates.

**Fig. 1.**
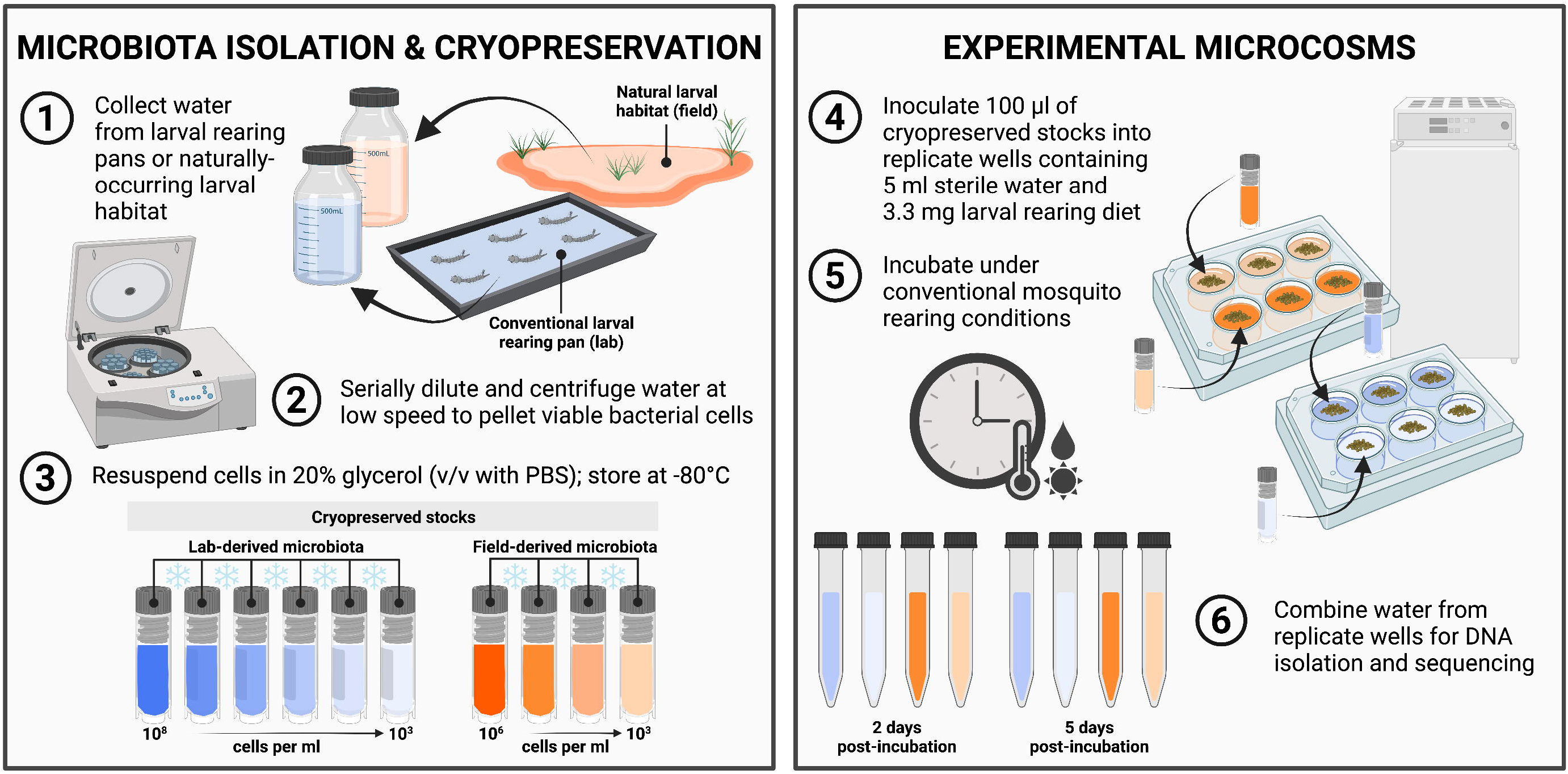
Overview of methodology used to isolate and cryopreserve microbiota and generate experimental microcosms. (*Left*) Isolation and cryopreservation of microbiota from conventional larval rearing pans of our laboratory colony of *A. aegypti* mosquitoes or a naturally occurring larval habitat in the field. Water from four replicate rearing pans containing larvae that had molted to the final (fourth) instar was collected from the lab, while four aliquots of water from the same larval habitat were collected from the field (1) prior to serial dilution and centrifugation at low speed to pellet any viable bacterial cells (2). The resulting cell pellets were then resuspended in 20% glycerol (v/v with PBS) and stored at −80°C to produce a total of 40 cryopreserved stocks at variable cell densities (~10^3^-10^8^ bacterial cells per ml) for use in downstream experiments (3). Aliquots of non-cryopreserved fresh water from each rearing pan or habitat water sample in (1) were also immediately centrifuged at high speed prior to removal of any supernatant and storage of pellets at −20°C for downstream DNA isolation and sequencing to characterize microbiota diversity. (*Right*) Generation of experimental microcosms colonized by standardized microbial communities. Aliquots of 100 ul from each cryopreserved stock were inoculated into replicate wells of 6-well culture plates containing 5 ml of sterile water and 3.3 mg of a standard larval diet sterilized by gamma irradiation (4). Replicate wells containing diet and 5 ml of non-cryopreserved fresh water from each rearing pan or habitat water sample in (1) served as unprocessed controls, while wells containing diet and sterile water only served as contamination controls. Culture plates were subsequently incubated under conventional mosquito rearing conditions (5) and water from replicate wells containing the same material was sampled 2- and 5-days post-incubation and combined (6) prior to centrifugation and storage as described above for downstream DNA isolation and sequencing. See “Methods” for more information. Created with BioRender.com.

### (b) Field-derived microbiota isolation and cryopreservation

We also isolated the microbiota present in a naturally occurring larval mosquito habitat identified and monitored annually by Public Health Madison & Dane County in Madison, WI USA. In brief, four 500-ml aliquots of water were collected in separate pre-sterilized containers and immediately transported on ice to the laboratory for processing as described above. A total of 16 cryopreserved stocks were produced at variable cell densities (~10^6^-10^3^ bacterial cells per ml). Cell density was estimated as described above.

### (c) Setup and inoculation of experimental microcosms

Sterile 6-well culture plates (Corning, Corning, NY, USA) were used for setting up microcosms (Fig. 1). Each microcosm consisted of a single well containing 3.3-mg of a standard mosquito larval rearing diet sterilized by gamma irradiation (8) and: (*i*) 5-ml of unprocessed water derived from either one of four larval rearing pans from our *A. aegypti* colony or one of four aliquots of water collected from a naturally occurring larval habitat in the field (unprocessed controls), (*ii*) 5-ml of sterile water plus 100-ul of material from a given cryopreserved stock generated from the same water, or (*iii*) 5-ml of deionized water sterilized by autoclaving in the laboratory (contamination control). A total of 300 microcosms were assayed (*n* = 3 wells per water source/treatment). Culture plates were incubated under conventional mosquito rearing conditions (27°C, 70% RH, 16-h light: 8-h dark photoperiod) and 5-ml of water from each well was sampled either 2- or 5-days post-incubation. Water samples collected from replicate wells containing the same water source/treatment were subsequently combined, centrifuged, and stored as described above for downstream DNA isolation and sequencing (Fig. 1).

### (d) Bacterial 16S rRNA library construction and sequencing

Total genomic DNA was isolated from a total of 108 samples using a standard phenol-chloroform extraction procedure (70) prior to one-step PCR amplification of the V4 region of the bacterial 16S rRNA gene using barcoded primers as described previously (71). PCR amplification was performed in 25-ul reactions containing ~10 ng of template DNA, 12.5-ul of 2X HotStart Ready Mix (KAPA Biosystems, Wilmington, MA, USA), and 5-pmol of each primer. No-template reactions as well as reactions using template from blank DNA extractions served as negative controls. Reaction conditions were: initial denaturation at 95°C for 3-min, followed by 25 cycles at 95°C for 30-sec, 58°C for 30-sec, and 72°C for 30-sec, and a final extension step at 72°C for 5-min. Products were visualized on 1% agarose gels and purified using a MagJET NGS Cleanup and Size Selection Kit (Thermo Fisher Scientific, Waltham, MA, USA). The resulting purified libraries were finally quantified using a Quantus fluorometer (Promega) and combined in equimolar amounts prior to paired-end sequencing (2 x 250-bp) on an Illumina MiSeq by the DNA Sequencing Facility at the University of Wisconsin-Madison (Madison, WI, USA).

### (e) Sequence analysis

De-multiplexed reads were processed using the DADA2 pipeline in QIIME 2-2021.2 (72,73). In brief, sequence reads were first filtered using DADA2’s recommended parameters. Filtered reads were then de-replicated and denoised using default parameters. After building the ASV table and removing chimeras, taxonomy was assigned using a Naïve Bayes classifier natively implemented in QIIME and pre-trained against the Silva reference database (138) (74). A phylogenetic tree was built using FastTree 2 (75) from a multiple sequence alignment made with the MAFFT alignment tool (76) against the Silva core reference alignment (74). All endpoint artifacts generated in QIIME were then exported, merged with metadata, and converted to a phyloseq object for further analysis in R (http://www.r-project.org/) (77). Rooting of the phylogenetic tree was performed in R using phyloseq and a decontamination procedure was implemented using a two-tiered approach implemented in the R package ‘decontam’ (78). In brief, DNA quantification values prior to library pooling in study samples, blank DNA extraction products, and PCR negative controls were used to generate a list of likely contaminant reads. Contaminant reads that were more prevalent in control samples than in study samples were then removed from the entire dataset, along with samples with fewer than 100 total reads and reads classified as ‘Archaea’, ‘Chloroplast’, or ‘mitochondria’ prior to downstream analyses.

Patterns of alpha diversity (as measured by Shannon’s H index and ASV richness) and beta diversity (as measured by the Bray-Curtis dissimilarity index) were analyzed using the R package ‘vegan’ (79). Differences in alpha and beta diversity between water samples collected from conventional larval rearing pans versus experimental microcosms were analyzed by Bonferroni-corrected pairwise Dunn’s tests to compare microcosm samples to their respective microbiota source (*e.g*., larval rearing pan). Differences in the proportion of rare and common taxa present in different microcosm samples were analyzed by Bonferroni-corrected Fisher’s exact tests. The significance of sample clustering by water source/treatment and time of sampling was analyzed by permutational multivariate analysis of variance (PERMANOVA), followed by *post-hoc* pairwise permutation tests for homogeneity of multivariate dispersions (PERMDISP). The differential abundance of ASVs between larval rearing pans and different experimental microcosm sample groups was analyzed based on a Wilcoxon rank sum test and Welch’s t test using the R package ALDEx2 (80–82). In order to identify ASVs significantly and systematically responding to cryopreservation, the ‘denom’ argument was set to “zero” to account for the complete loss of some taxa in samples derived from microcosms containing cryopreserved microbiota while allowing for all taxa present in at least one rearing pan or habitat water sample to be included in the analysis. To determine the effect of cryopreservation on bacterial populations, the effect size was calculated, which is the median of the ratio of the between group difference and the larger of the variances within groups. The Benjamini-Hochberg-Yekutieli procedure was then used to account for multiple testing, and corrected values were expressed as false discovery rates (FDR) (83). Finally, statistically significant differences in the average relative abundance of reads from shared taxa among replicates of different sample groups were assessed using Bonferroni-corrected pairwise Dunn’s tests followed by pairwise Fligner-Killeen tests to assess the homogeneity of variances. The same tests were then used to compare patterns of alpha and beta diversity among replicates of the same sample groups.

## 3. Results

### (a) Impact of cryopreservation on recovery of bacteria from laboratory-derived larval rearing pans

We initially sought to develop an approach to isolate and cryopreserve microbiota from conventional larval rearing pans in the laboratory for recapitulation in experimental microcosms. Multiplex sequencing of 16S rRNA gene amplicons for the resulting rearing pan- and microcosm-derived water samples generated a total of 2,669,979 quality-filtered reads with a median sequencing depth of 44,707 reads per sample (Supplementary Table 1). An unusually low number of reads (<100) were obtained for two samples, which were removed from subsequent analyses (Supplementary Table 1). Rarefaction curves for the remaining samples saturated at ~500 sequences, indicating that most (if not all) bacteria in each sample were captured (Supplementary Fig. 1).

We identified 94 ASVs across the conventional larval rearing pans we sampled. However, the vast majority (>92%) of reads from these samples were assigned to ASVs belonging to one of eight bacterial families within the following phyla/classes: Alphaproteobacteria (Azospirillaceae, Rhizobiaceae, Sphingomonadaceae), Betaproteobacteria (Comamonadaceae), Gammaproteobacteria (Enterobacteriaceae, Moraxellaceae), and Bacteroidetes (Sphingobacteriaceae, Weeksellaceae) (Fig. 2A). A total of 85 of the 94 ASVs found in rearing pans, representing >99% of all rearing pan sequences, were also detected in the experimental microcosms we set-up and sampled (Fig. 2B), although recovery varied with respect to the time of microcosm sampling, the cell density of the cryopreserved stock used to generate a given microcosm, and how common a given ASV was across the rearing pans we sampled and sequenced (Supplementary Fig. 2). Cryopreservation had the greatest impact on recovery of rare ASVs (*i.e*., those with a maximum relative abundance ≤1% in rearing pans), with significantly fewer rare ASVs being recovered in experimental microcosms generated using cryopreserved stocks of lower cell densities (10^4^, 10^5^), even at 5 days post-incubation (Supplementary Fig. 2). However, there were no significant differences in recovery of more common ASVs (*i.e*., those with a minimum relative abundance >1% in rearing pans) between any of the microcosms we generated (Supplementary Fig. 2).

**Fig. 2.**
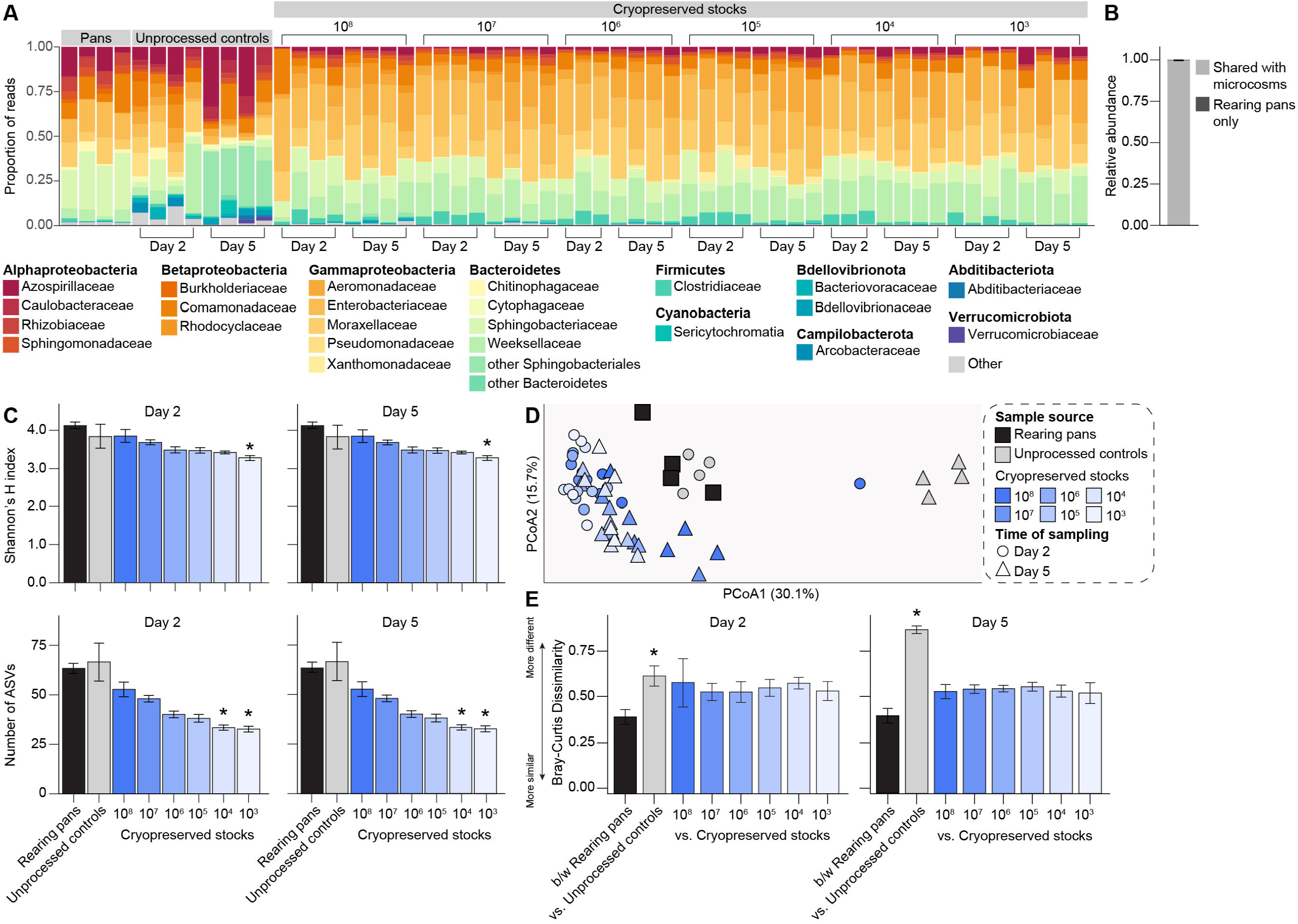
Bacterial diversity in laboratory larval rearing pans and experimental microcosms generated using lab-derived microbiota. (A) Relative abundance of bacterial families in water sampled from: (*i*) conventional rearing pans containing fourth instar larvae from our standard *A. aegypti* laboratory colony, (*ii*) experimental microcosms containing unprocessed water from the same rearing pans, or (*iii*) experimental microcosms containing water plus material from a given cryopreserved stock. Each bar presents the proportion of sequencing reads assigned to a given bacterial family. Low abundance families (<1%) are represented by the ‘Other’ category. (B) Average relative abundance of rearing pan microbiota shared with experimental microcosms. (C) Alpha diversity of rearing pans and experimental microcosms, as measured by Shannon’s H index (top) and ASV richness (bottom). Mean values ± standard errors are shown. Asterisks (*) indicate significant differences between a given group of experimental microcosm samples relative to rearing pan samples (Dunn’s test with Bonferroni correction, *P* < 0.01). (D) Principal coordinates analysis using the Bray-Curtis dissimilarity index. Symbols are colored by sample source (rearing pans, black; experimental microcosms containing unprocessed water, grey; experimental microcosms containing water plus material from a given cryopreserved stock, blue). Time of sampling (Day 2 or Day 5) of experimental microcosms is designated by symbol shape. (E) Average Bray-Curtis dissimilarity between (b/w) rearing pans versus between a given rearing pan and group of experimental microcosm samples. Mean values ± standard errors are shown. Asterisks (*) indicate comparisons for which the average dissimilarity between a given rearing pan and group of experimental microcosm samples was significantly higher than that expected as a result of the microbiota isolation and cryopreservation procedure itself (*i.e*., between rearing pans) (Dunn’s test with Bonferroni correction, *P* < 0.05).

### (b) Recapitulation of microbial diversity in experimental microcosms

Experimental microcosms generated using material from cryopreserved stocks also generally contained bacterial communities that did not significantly differ in alpha diversity from the communities present in the rearing pans we initially sampled, with the only exceptions again being those generated using cryopreserved stocks of lower cell densities (10^4^, 10^5^) (Fig. 2C; Supplementary Table 1). Further, while a principal coordinates analysis using the Bray-Curtis dissimilarity index identified significant clustering by sample source (*i.e*., rearing pans vs. experimental microcosms containing unprocessed water or water plus material from cryopreserved stocks; PERMANOVA, *P* = 0.001) (Fig. 2D), *post-hoc* pairwise permutation tests for homogeneity of multivariate dispersions revealed significantly higher dispersion values amongst experimental microcosms containing unprocessed water than amongst microcosms containing water plus material from cryopreserved stocks or the larval rearing pans (PERMDISP, *P* = 0.001). Differences in beta diversity, measured as average Bray-Curtis dissimilarity, were also overall higher between rearing pans and unprocessed controls than between rearing pans and experimental microcosms generated using material from cryopreserved stocks, regardless of cell density (Fig. 2E).

### (c) Changes in the relative abundance of specific taxa in response to cryopreservation

Finally, we performed ALDEx2 tests to identify rearing pan ASVs that significantly changed in relative abundance in response to cryopreservation and/or re-culturing under conventional mosquito rearing conditions (Fig. 3; Supplementary Figs. 3 & 4). These tests revealed that 30 of the 94 ASVs found in rearing pans, classified as members of one of four bacterial phyla (Actinobacteria, Bacteroidetes, Firmicutes, and Proteobacteria), showed significantly different relative abundances in experimental microcosms containing unprocessed water and/or water plus material from cryopreserved stocks (Fig. 3; Supplementary Figs. 3 & 4). However, only 10 of these ASVs, representing <7% of all rearing pan sequences, were specifically negative affected (*i.e*., decreased in abundance) in response to cryopreservation (Fig. 3; Supplementary Figs. 3 & 4). The vast majority (>94%) of rearing pan ASVs recovered in microcosms containing material from cryopreserved stocks also persisted over time, with only four ASVs showing significantly lower relative abundances in samples collected on Day 5 when compared to those collected on Day 2 post-incubation (ALDEx2, *P* < 0.05; FDR, *P* < 0.05).

**Fig. 3.**
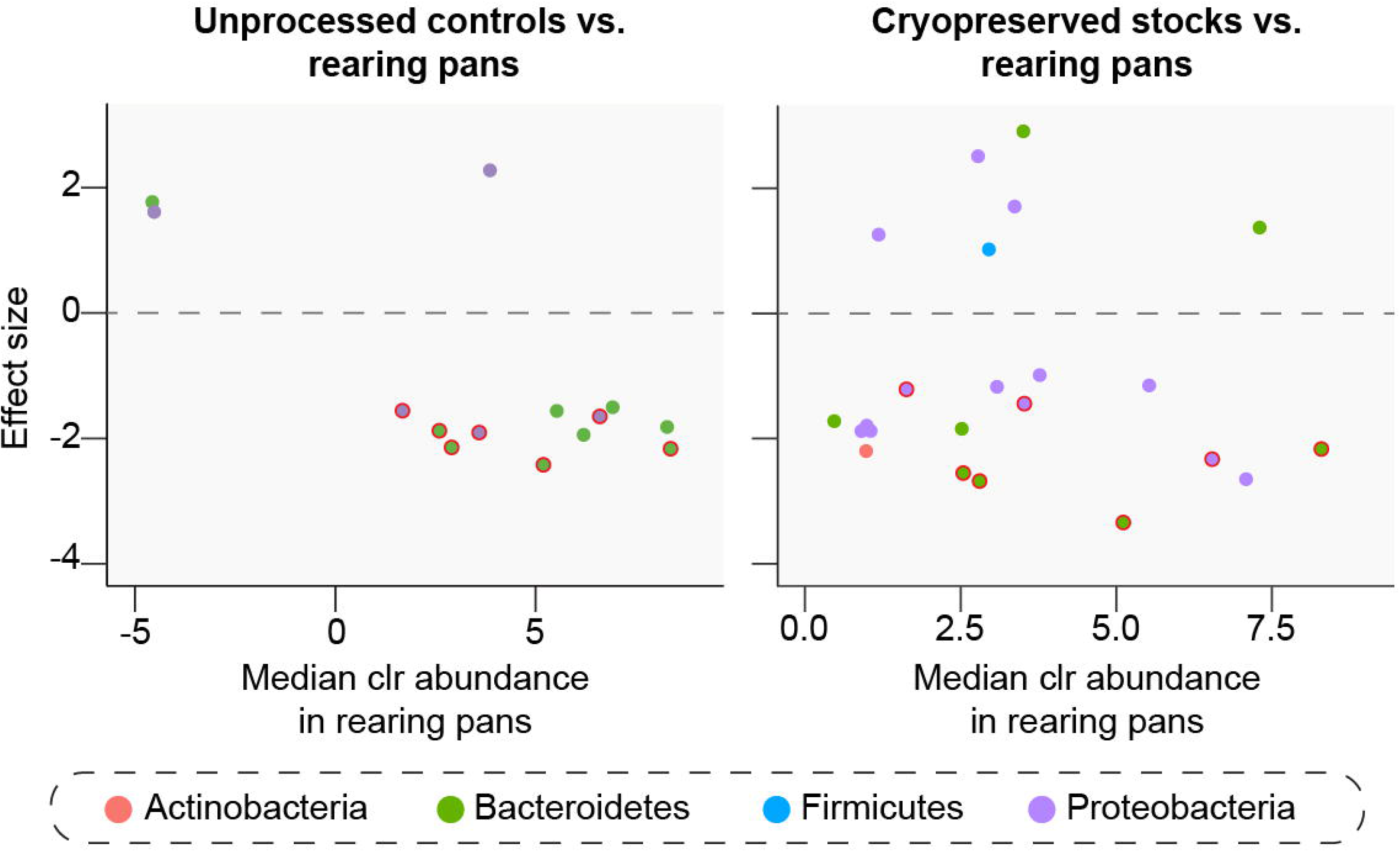
ASVs that significantly varied in abundance between larval rearing pans and experimental microcosms containing unprocessed water (*left*) or water plus material from cryopreserved stocks (*right*) (ALDEx2, *P* < 0.05; FDR, *P* < 0.05). Plots show the median clr value for each ASV across larval rearing pans (*x* axis) and the effect of re-culturing with or without cryopreservation under conventional mosquito rearing conditions (*y* axis). An effect size < 0 indicates that the ASV abundance significantly decreased between rearing pans and a given group of experimental microcosm samples, and an effect size of > 0 indicates that the ASV abundance significantly increased. ASVs are colored by phylum (see legend). ASVs with a bold red outline were differentially abundant in both experimental microcosms containing unprocessed water and experimental microcosms containing water plus material from cryopreserved stocks.

### (d) Validation of isolation and cryopreservation approaches using field-derived microbiota

Next we sought to validate the use of cryopreservation approaches developed above to generate experimental microcosms colonized by microbiota derived from a naturally occurring mosquito larval habitat in the field. Multiplex sequencing of 16S rRNA gene amplicons for the resulting habitat- and microcosm-derived water samples generated a total of 2,111,045 quality-filtered reads with a median sequencing depth of 43,541 reads per sample (Supplementary Table 2), with rarefaction curves for most samples saturating at ~3,000 sequences (Supplementary Fig. 5).

As expected, we identified a substantially higher diversity of bacteria (993 ASVs) across the field-derived water samples as compared to those derived from conventional larval rearing pans in the laboratory, with ~42% of reads being assigned to ASVs belonging to the eight bacterial families dominating communities in rearing pans and the remaining ~58% being assigned to ASVs belonging to one of 306 other families within 187 bacterial orders and 34 phyla/classes (Fig. 4A). A substantially higher diversity of bacteria (182 ASVs) was also detected in the experimental microcosms we set-up and sampled, although overall recovery was significantly lower with respect to the relative abundance of habitat ASVs that were shared with experimental microcosms (Fig. 4B) and impacts on alpha diversity were more marked in microcosms generated using cryopreserved stocks of higher cell densities (Fig. 4C; Supplementary Table 2) than previously observed for cryopreserved stocks of lab-derived microbiota (Fig. 2C; Supplementary Table 1). Differences in beta diversity, measured as Bray-Curtis dissimilarity, were also overall higher between habitat samples and microcosms containing material from cryopreserved stocks (Fig. 4D,E) than previously observed for lab-derived bacterial communities (Fig. 2D,E), although no dispersion effects were detected (PERMDISP, *P* = 0.426).

**Fig. 4.**
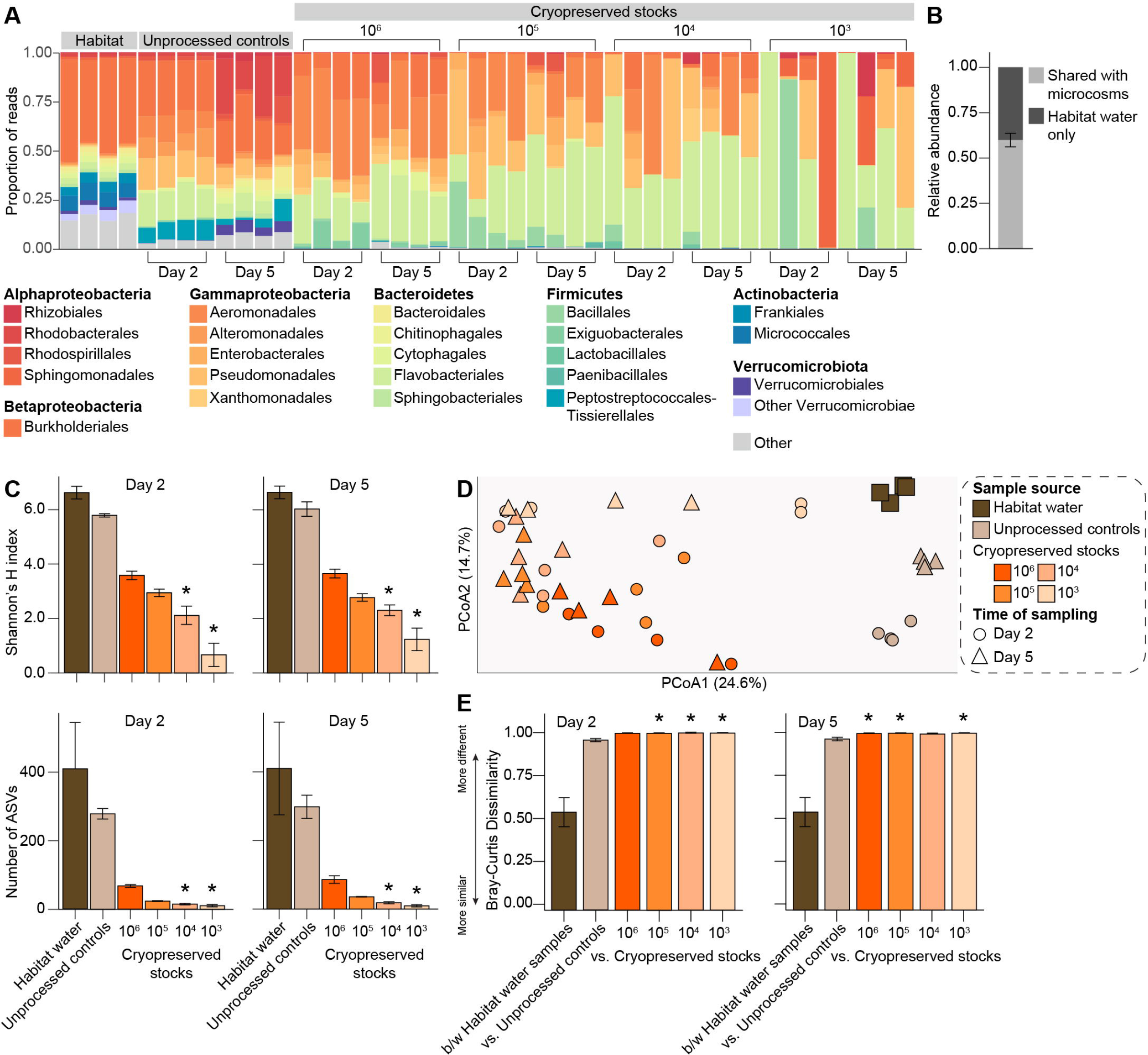
Bacterial diversity in a natural larval habitat and experimental microcosms generated using field-derived microbiota. (A) Relative abundance of bacterial families in water sampled from: (*i*) a naturally occurring larval mosquito habitat in the field, (*ii*) experimental microcosms containing unprocessed water from the same habitat, or (*iii*) experimental microcosms containing water plus material from a given cryopreserved stock. Each bar presents the proportion of sequencing reads assigned to a given bacterial family. Low abundance families (<1%) are represented by the ‘Other’ category. (B) Average relative abundance of habitat water microbiota shared with experimental microcosms. (C) Alpha diversity of habitat water samples and experimental microcosms, as measured by Shannon’s H index (top) and ASV richness (bottom). Mean values ± standard errors are shown. Asterisks (*) indicate significant differences between a given group of experimental microcosm samples relative to habitat water samples (Dunn’s test with Bonferroni correction, *P* < 0.05). (D) Principal coordinates analysis using the Bray-Curtis dissimilarity index. Symbols are colored by sample source (habitat water, brown; experimental microcosms containing unprocessed water, tan; experimental microcosms containing water plus material from a given cryopreserved stock, orange). Time of sampling (Day 2 or Day 5) of experimental microcosms is designated by symbol shape. (E) Average Bray-Curtis dissimilarity between (b/w) habitat water samples versus between a given habitat water sample and group of experimental microcosm samples. Mean values ± standard errors are shown. Asterisks (*) indicate comparisons for which the average dissimilarity between a given habitat water sample and group of experimental microcosm samples was significantly higher than that expected as a result of the microbiota isolation and cryopreservation procedure itself (*i.e*., between habitat water samples) (Dunn’s test with Bonferroni correction, *P* < 0.05).

ALDEx2 tests further supported a greater influence of our isolation and cryopreservation approaches on the recovery and relative abundance of specific taxa in experimental microcosms containing field-derived microbiota as compared to lab-derived microbiota, while also strongly suggesting that shifts in diversity were more likely due to the inability of many taxa to persist under conventional mosquito rearing conditions and/or utilize standard larval mosquito diets rather than the cryopreservation process itself. A total of 110 of the 993 ASVs found in the habitat water samples we sequenced showing significantly different relative abundances in experimental microcosms, but only 12 of these ASVs, representing <6% of all habitat water sequences, were specifically negatively affected in response to cryopreservation (Fig. 5; Supplementary Figs. 6 & 7). In contrast, 87 ASVs, representing ~68% of all habitat water sequences, were negatively affected in both experimental microcosms containing unprocessed water and microcosms containing material from cryopreserved stocks (Fig. 5; Supplementary Figs. 6 & 7). However, similar to our ALDEx2 results for microcosms containing lab-derived microbiota, all (100%) of the habitat ASVs recovered in microcosms generated using cryopreserved stocks containing field-derived microbiota persisted over time, with no ASVs showing significantly lower relative abundances between samples collected on Day 5 and Day 2 post-incubation (ALDEx2, *P* > 0.05).

**Fig. 5.**
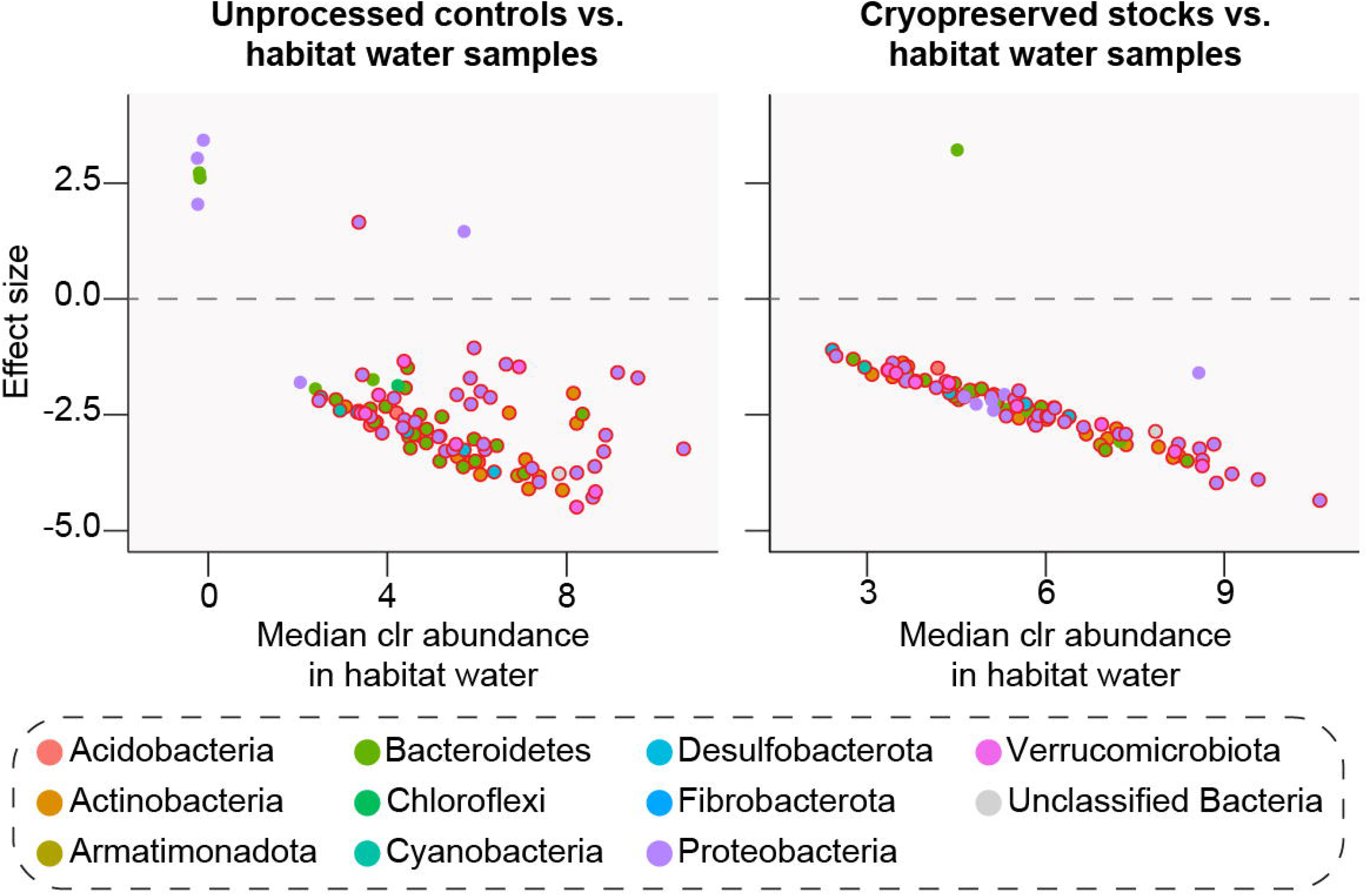
ASVs that significantly varied in abundance between habitat water samples and experimental microcosms containing unprocessed water (*left*) or water plus material from cryopreserved stocks (*right*) (ALDEx2, *P* < 0.05; FDR, *P* < 0.05). Plots show the median clr value for each ASV across habitat water samples (*x* axis) and the effect of re-culturing with or without cryopreservation under conventional mosquito rearing conditions (*y* axis). An effect size < 0 indicates that the ASV abundance significantly decreased between habitat water samples and a given group of experimental microcosm samples, and an effect size of > 0 indicates that the ASV abundance increased. ASVs are colored by phylum (see legend). ASVs with a bold red outline were differentially abundant in both experimental microcosms containing unprocessed water and experimental microcosms containing water plus material from cryopreserved stocks.

### (e) Reproducibility of microbial communities in experimental microcosms

While our isolation and cryopreservation approaches resulted in the loss of some lab- and field-derived bacterial taxa and concurrent changes in the total alpha and beta diversity of microbiota within experimental microcosms as compared to the larval rearing pans and habitat water we originally sampled, we observed little variation among the bacterial communities present within replicate microcosms generated using the same cryopreserved stock, with the only exceptions being those generated using stocks of lower cell densities (Fig. 6). On average, replicate microcosms shared 30 taxa, which accounted for ~96% of their total reads– consistent with the amount of bacteria shared among larval rearing pans in the laboratory (Fig. 6A). Bacteria introduced into replicate microcosms also assembled into communities that overall exhibited similar patterns of inter-replicate alpha and beta diversity as those observed in rearing pans under conventional rearing conditions (Fig. 6B,C), indicating that we were successful in producing experimental microcosms colonized by standardized microbial communities.

**Fig. 6.**
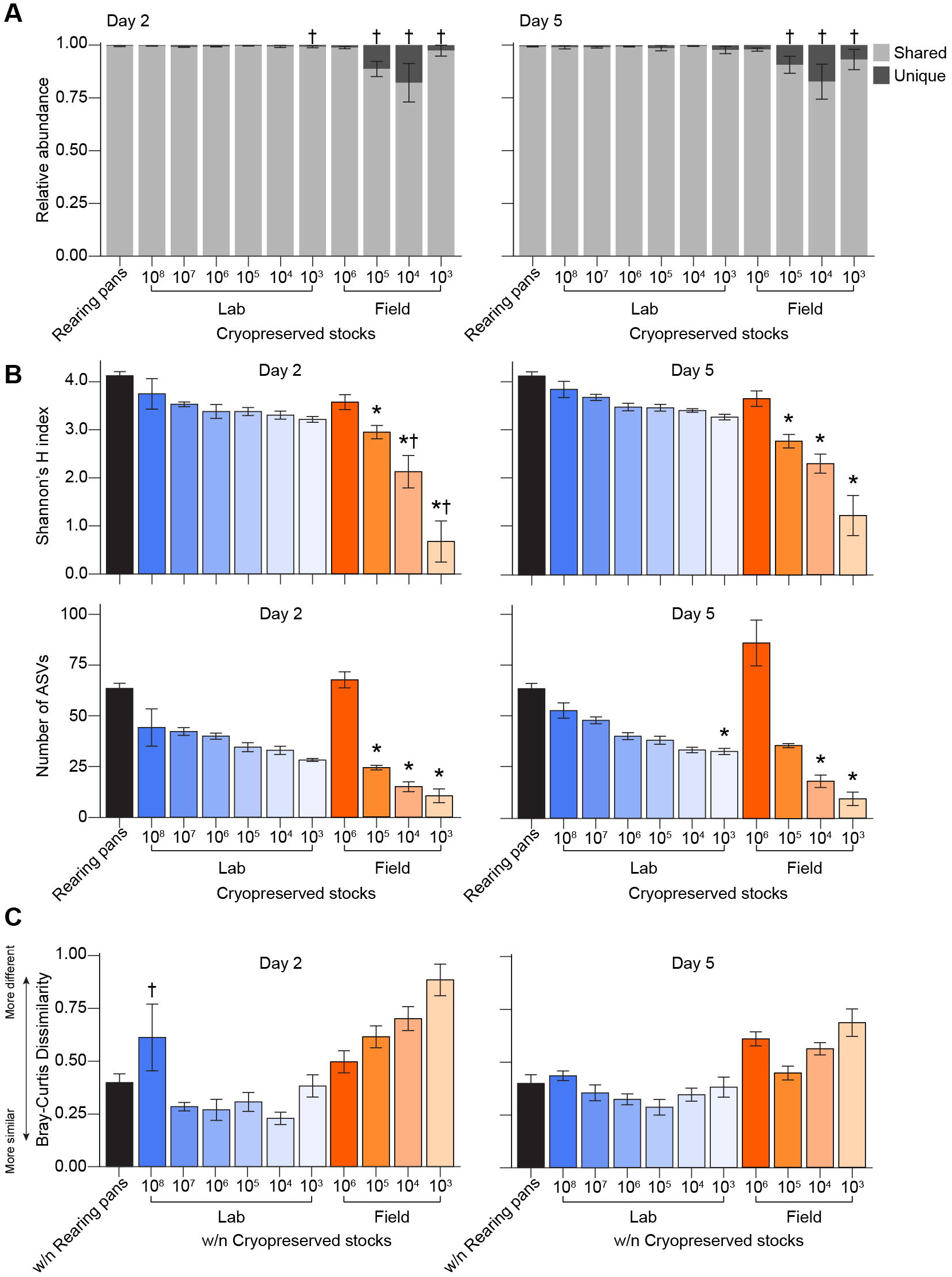
Reproducibility of bacterial communities in replicate microcosms. (A) Average relative abundance of shared microbiota among rearing pan samples and replicates of a given group of experimental microcosms. Daggers (†) indicate comparisons for which the variance of a given group of experimental microcosm samples was significantly higher than that of rearing pan samples (Fligner-Killeen test, *P* < 0.05). No significant differences in averages were detected between rearing pan samples and any experimental microcosm sample group (Dunn’s test with Bonferroni correction, *P* > 0.05). (B) Alpha diversity of rearing pans and experimental microcosms generated using cryopreserved stocks of lab- and field-derived microbiota, as measured by Shannon’s H index (top) and ASV richness (bottom). Mean values ± standard errors are shown. Asterisks (*) indicate significant differences between a given group of microcosm samples relative to rearing pan samples (Dunn’s test with Bonferroni correction, *P* < 0.05). Daggers (†) indicate comparisons for which the variance of a given group of experimental microcosm samples was significantly higher than that of rearing pan samples (Fligner-Killeen test, *P* < 0.05). (C) Average Bray-Curtis dissimilarity within (w/n) rearing pans versus within a given group of experimental microcosm samples. Mean values ± standard errors are shown. Daggers (†) indicate comparisons for which the variance of a given group of experimental microcosm samples was significantly higher than that of rearing pan samples (Fligner-Killeen test, *P* < 0.01). No significant differences in averages were detected between rearing pan samples and any experimental microcosm sample group (Dunn’s test with Bonferroni correction, *P* > 0.05).

## 4. Discussion

Mosquitoes live in close association with bacteria and other microorganisms that shape their ability to transmit pathogens (1). However, the immense diversity and variability of the microbiota within and between different populations of mosquitoes in the laboratory and field have made studying mosquito-microbiome interactions–and identifying bacteria that reduce the vectorial capacity of mosquitoes–a formidable challenge (1). As in other animals, numerous factors have the potential to shape variation in mosquito microbiota and therefore variation in mosquito phenotypes, including the microbiota present in the aquatic environment in which larvae develop, environmental conditions (e.g., diet, temperature), and host genetics (2–16,51,52,57,58,60–62,84–92). However, while experiments with mosquitoes are commonly conducted using genetically identical individuals under highly controlled environmental conditions, tools to standardize the microbiota present in the larval rearing environment are comparatively limited (8,55,93–96).

In this study, we first developed an approach to isolate and cryopreserve microbiota from conventional larval rearing pans in the laboratory for recapitulation in experimental microcosms. High-throughput sequencing of 16S rRNA gene amplicons from water collected from four replicate rearing pans from our laboratory colony of *A. aegypti* revealed a bacterial community comprised of ~100 unique bacterial taxa and dominated by members of the Proteobacteria, Bacteroidetes, and Firmicutes, consistent with sequencing studies of other laboratory colonies of mosquitoes (2–4,6–16,45,47,57,60,92,97). The vast majority of this bacterial diversity was also recovered in both experimental microcosms generated using non-cryopreserved fresh material and experimental microcosms generated using cryopreserved material from the same rearing pans, regardless of stock cell density. The only exceptions were a handful of representatives of the class Alphaproteobacteria, members of the orders Burkholderiales, Chitinophagales, and Sphingobacteriales, as well as one very rare ASV belonging to the genus *Paenarthrobacter*, which were significantly reduced or absent in experimental microcosms and may represent taxa that are not readily amenable to centrifugation and/or cryopreservation using our methodology.

We next validated our isolation and cryopreservation approach using microbiota derived from water collected from a naturally occurring larval habitat in the field. As expected, the bacterial community present in this habitat was comprised of a substantially higher diversity of ~1,000 unique taxa, although these taxa were dominated by members of the same phyla (Proteobacteria, Bacteroidetes, Firmicutes, Actinobacteria) detected in laboratory rearing pans and commonly detected in field-collected mosquitoes (5,51,52,56–62,84–91,97–102). Recovery of field-derived taxa in experimental microcosms generated using cryopreserved material from the same habitat was also overall much lower than what we observed in microcosms generated using cryopreserved stocks of lab-derived microbiota, although the taxa lost or significantly reduced in microcosms generated using cryopreserved material were consistent with those lost or significantly reduced in microcosms generated using fresh material–strongly suggesting that the observed shifts in alpha and beta diversity in microcosms were not the result of cryopreservation but rather the inability of many field-derived taxa to thrive under conventional mosquito rearing conditions in the laboratory. This, combined with our results using lab-derived microbiota, strongly suggests that our isolation and cryopreservation procedure was sufficient to conserve most bacteria. Nevertheless, future studies could compare the results here to results obtained using different long-term preservation approaches, including periodic subculturing, drying, freeze-drying, and cryopreservation with different cryoprotectants at variable concentrations (63,65,66,103–108). Future work will also be necessary to confirm that the patterns observed here are the same for experimental microcosms generated using material from cryopreserved stocks stored for longer periods of time (109) and to formally assess the impact of long-term cryopreservation and resuscitation on cell viability and functional stability (110,111). Additionally, it would be important to determine that the cryopreservation approach is appropriate for microbiomes collected from diverse environments given the variation between different habitats (5,57,58,62,84–86,89–91).

The cell density of cryopreserved stocks also had a much larger impact on the recovery and persistence of field-derived taxa, with microcosms generated using stocks of lower cell densities exhibiting more variable bacterial communities that were overall lower in complexity that communities in microcosms generated using stocks of higher cell densities. This is consistent with results showing that small sampling volumes from aquatic and other high-complexity environments like soil often fail to adequately capture bacterial diversity (112–114), owing to microscale heterogeneity and bottleneck effects during population sub-sampling that leads to the stochastic proliferation of some taxa and the loss of others (115–122).

Overall, experimental microcosms generated using cryopreserved stocks containing lab-derived microbiota exhibited levels of reproducibility that were the same or better than those present under conventional rearing conditions in the laboratory, highlighting the value of our approach for intra-and inter-lab studies of mosquitoes in the presence of standardized microbial communities. Similar levels of reproducibility could also be replicated in microcosms generated using cryopreserved stocks of field-derived microbiota, although reproducibility significantly decreased with decreasing inoculum size–consistent with our previous observation that variability among the bacterial communities present within replicate microcosms was significantly higher in those generated using cryopreserved stocks of lower cell densities. Future studies are warranted to establish best practices for field habitat sampling, though our results at minimum suggest that higher-volume samples are preferable, and that sampling should be avoided in situations where bacterial density is expected to be low (*e.g*., immediately after a rainfall). Defined diets that support mosquito growth and development could also be developed to improve the reproducibility of field bacterial diversity between replicate microcosms, including the maintenance of rare taxa important for community assembly and stability (123,124).

Owing to their ability to impact numerous components of mosquito vectorial capacity, there is a growing interest in exploiting microbes for mosquito-borne disease control. For example, bacteria that naturally colonize the mosquito gut could be genetically modified to produce effector molecules that alter the mosquito’s ability to become infected with and transmit pathogens, or that reduce mosquito fecundity or lifespan (*i.e*., paratransgenesis). Unmodified bacteria that naturally inhibit pathogen colonization or mosquito fitness could also be disseminated to mosquito populations. The identification of suitable microbial candidates for pathogen or mosquito control will require a comprehensive understanding of the factors that influence the acquisition, maintenance, and transmission of mosquito microbiota and the mechanisms that underlie how individual microbial species and assemblages impact mosquito vectorial capacity. However, the dearth of tools to manipulate mosquito microbiota, including microbiota derived from naturally occurring mosquito habitats in the field, has greatly slowed progress in this area. In this way, the methods reported here not only provide a critical first step toward the standardization of microbial inputs in mosquito studies, but also provide a critical first step toward the identification of taxonomic and functional profiles of bacteria associated with phenotypic traits of interest in mosquitoes. For example, a defined microbiome could be universally adopted by the community to conduct vector competence assays in the absence of confounding effects due to microbiota variation. Libraries of cryopreserved microbiota could also be screened to identify bacteria that improve or reduce mosquito fitness and/or pathogen susceptibility, or to predict the success of individual microbial candidates under variable microbial conditions. Similarly, the results reported here strongly support that our methods could immediately be leveraged to expand studies of mosquito-microbiota interactions to include microbial genotypes derived from the field, and thereby conduct lab-based mosquito experiments with a field-relevant microbiome.

## Acknowledgments

We thank Lyric Bartholomay and Kathy Vaccaro for assistance with maintaining the *A. aegypti* colony used in this study. We also thank Sarah Hilby, Abby Cook, and John Hausbeck from Public Health Madison & Dane County for assistance with field sampling, as well as Candace Davison and the Radiation Science & Engineering Center at The Pennsylvania State University for assistance in preparing irradiated diet for use in our experiments.

## Data Availability

Raw Illumina reads are available in the NCBI Sequence Read Archive (https://www.ncbi.nlm.nih.gov/sra) under BioProject ID PRJNA856768. Input files for the QIIME pipeline as well as raw data files and R code for statistical analyses have been deposited in the Dryad Digital Repository under doi:10.5061/dryad.dfn2z354z.

## Author Contributions

SYZ, GLH, and KLC conceived and designed the experiments. SYZ performed the experiments. SYZ and KLC carried out the data analysis. SYZ and KLC wrote the initial manuscript, and GLH contributed to revisions.

