## Supplementary Information for "Cryopreservation of mosquito microbiota for use in microcosm experiments"

**Contents:**

Supplementary Figure 1 | Rarefaction data from Illumina sequencing of water from conventional larval rearing pans in the laboratory and resulting experimental microcosms

Supplementary Figure 2 | Proportion of rare and abundant ASVs found in at least one larval rearing pan that were detected in experimental microcosms

Supplementary Figure 3 | Differentially abundant ASVs in water collected from conventional larval rearing pans in the laboratory and experimental microcosms containing unprocessed pan water

Supplementary Figure 4 | Differentially abundant ASVs in water collected from conventional larval rearing pans in the laboratory and experimental microcosms containing water plus material from cryopreserved stocks

Supplementary Figure 5 | Rarefaction data from Illumina sequencing of water samples from a naturally occurring mosquito larval habitat in the field and resulting experimental microcosms

Supplementary Figure 6 | Differentially abundant ASVs in water samples collected from a naturally occurring mosquito larval habitat in the field and experimental microcosms containing unprocessed habitat water

Supplementary Figure 7 | Differentially abundant ASVs in water samples collected from a naturally occurring mosquito larval habitat in the field and experimental microcosms containing water plus material from cryopreserved stocks

Supplementary Table 1 | Sequencing and diversity statistics for 16S rRNA gene amplicon libraries prepared from water collected from conventional larval rearing pans in the laboratory and resulting experimental microcosms

Supplementary Table 2 | Sequencing and diversity statistics for 16S rRNA gene amplicon libraries prepared from water collected from a naturally occurring mosquito larval habitat in the field and resulting experimental microcosms

Supplementary Figure 1

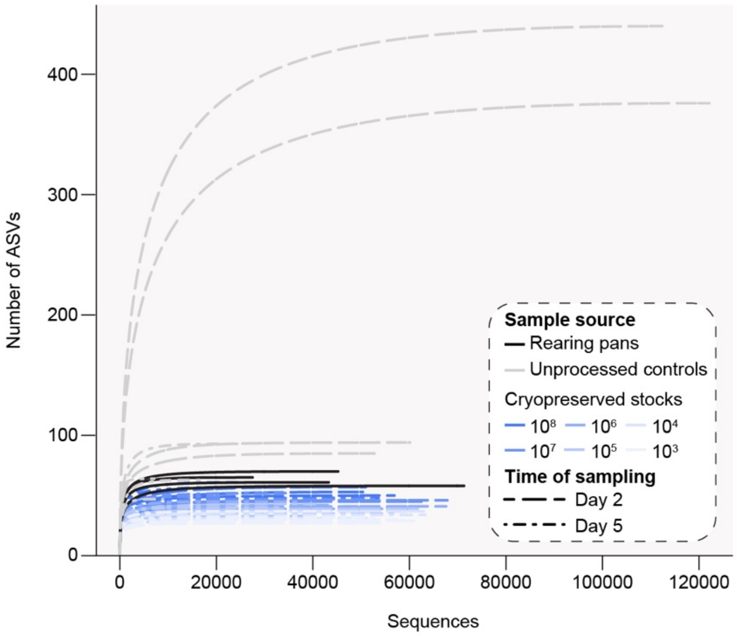


**Supplementary Fig. 1**. Rarefaction data from Illumina sequencing of water from conventional larval rearing pans in the laboratory and resulting experimental microcosms. Reads from each water library were sampled starting at 1 sequence per step and increased in increments of 100 until the total number of reads per sample was reached. Lines are colored by sample source (rearing pans, black; experimental microcosms containing unprocessed water, grey; experimental microcosms containing water plus material from a given cryopreserved stock, blue). Time of sampling of experimental microcosms is designated by line type (Day 2, long-dash; Day 5, dot-dash).

Supplementary Figure 2

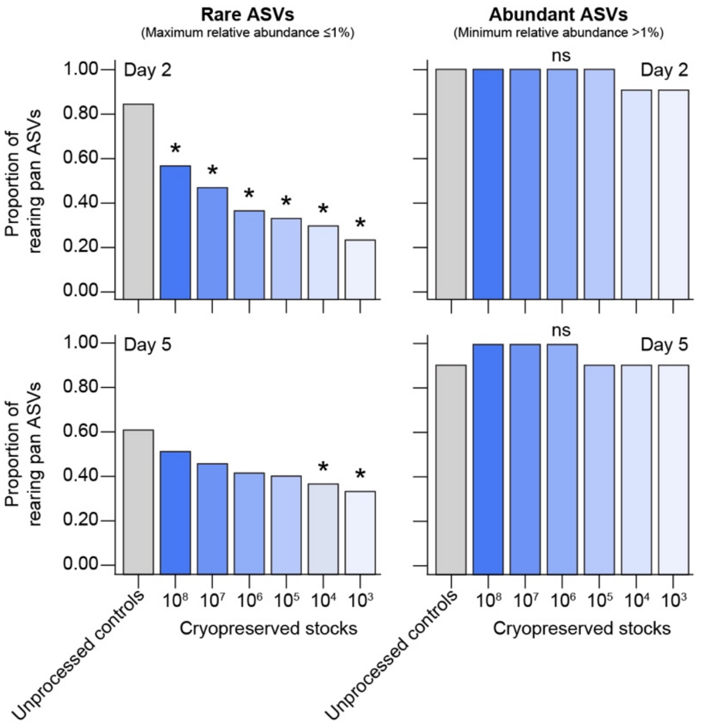


**Supplementary Fig. 2**. Proportion of rare (left) and abundant(right) ASVs found in at least one larval rearing pan that were detected in experimental microcosms containing unprocessed water or water plus material from a given cryopreserved stock. An ASV was considered "rare" if it had a maximum relative abundance ≤1% across the four larval rearing pans we sampled, while ASVs with a minimum relative abundance >1% were considered "abundant". Asterisks (*) indicate significant differences between experimental microcosms generated using cryopreserved stocks relative to unprocessed controls as determined by paired Fisher's exact tests with Bonferroni correction ($P < 0.05$).

Supplementary Figure 3

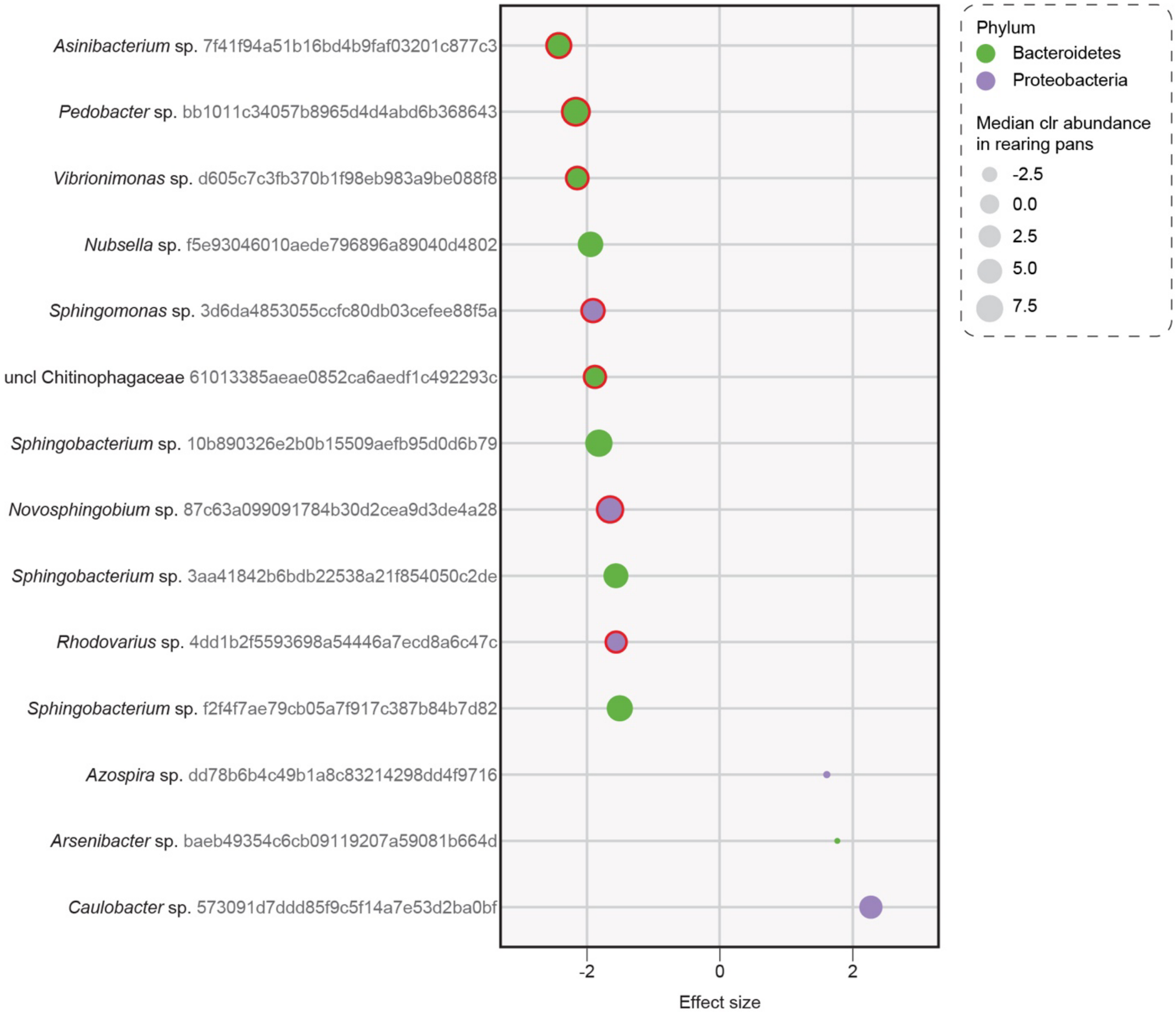


**Supplementary Fig. 3**. Differentially abundant ASVs in water collected from conventional larval rearing pans in the laboratory and experimental microcosms containing unprocessed pan water, identified through ALDEx2 testing (ALDEx2, $P < 0.05$; FDR, $P < 0.05$). Each ASV is presented with its lowest annotated taxonomic rank (to genus level) together with its ASV ID. The ASVs are color-coded according to the phyla they belong to and plotted according to their effect size, calculated as the levels in samples from experimental microcosms relative to levels in samples from larval rearing pans. Dot sizes correspond to the median clr abundance value for each ASV across rearing pan samples. Dots with a bold red outline represent ASVs that were differentially abundant in both experimental microcosms containing unprocessed water and experimental microcosms containing water plus material from cryopreserved stocks (see Supplementary Fig. 4).

### Supplementary Figure 4

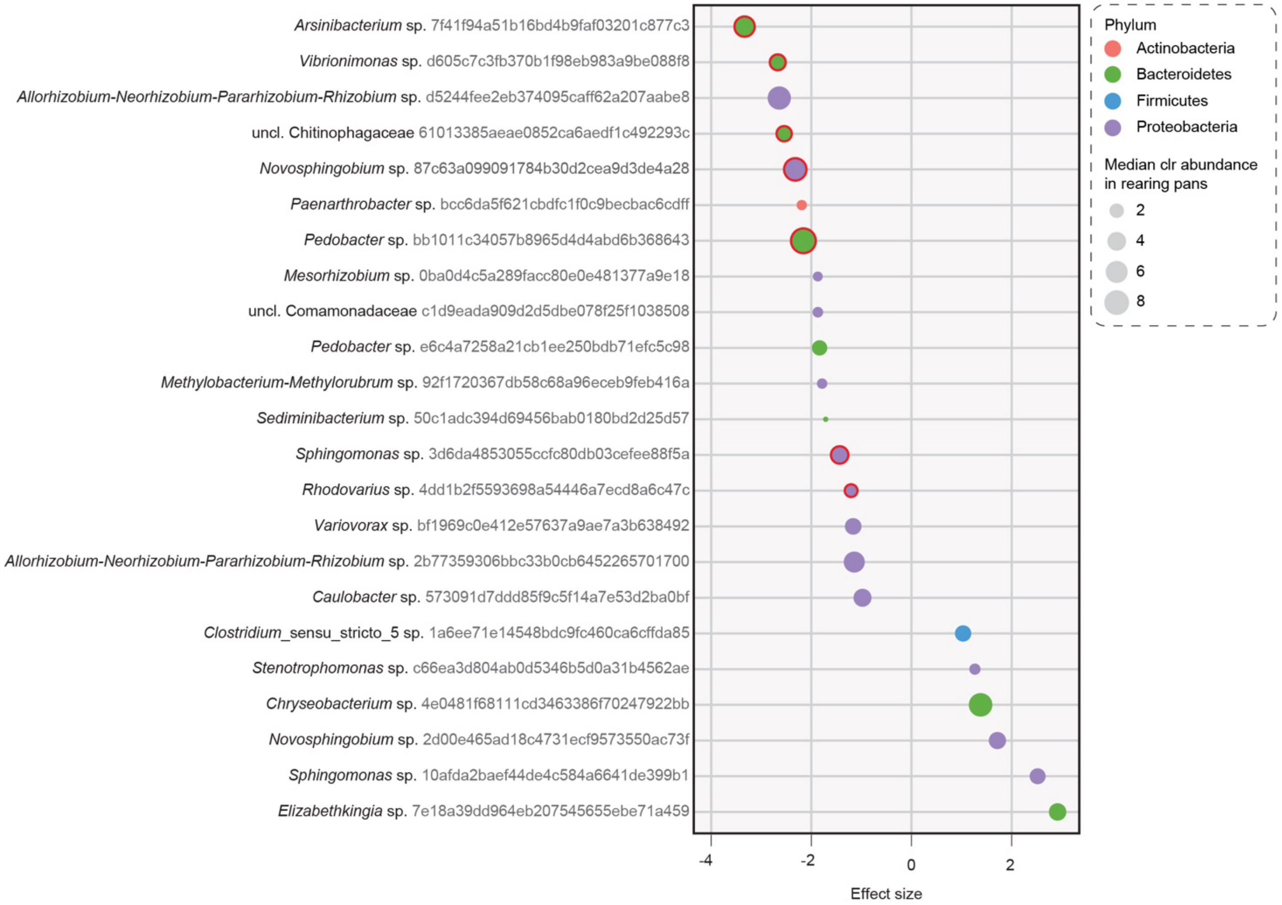


**Supplementary Fig. 4**. Differentially abundant ASVs in water collected from conventional larval rearing pans in the laboratory and experimental microcosms containing water plus material from cryopreserved stocks, identified through ALDEx2 testing (ALDEx2, P < 0.05; FDR, P < 0.05). Each ASV is presented with its lowest annotated taxonomic rank (to genus level) together with its ASV ID. The ASVs are color-coded according to the phyla they belong to and plotted according to their effect size, calculated as the levels in samples from experimental microcosms relative to levels in samples from larval rearing pans. Dot sizes correspond to the median clr abundance value for each ASV across rearing pan samples. Dots with a bold red outline represent ASVs that were differentially abundant in both experimental microcosms containing water plus material from cryopreserved stocks and experimental microcosms containing unprocessed water (see Supplementary Fig. 3).

Supplementary Figure 5

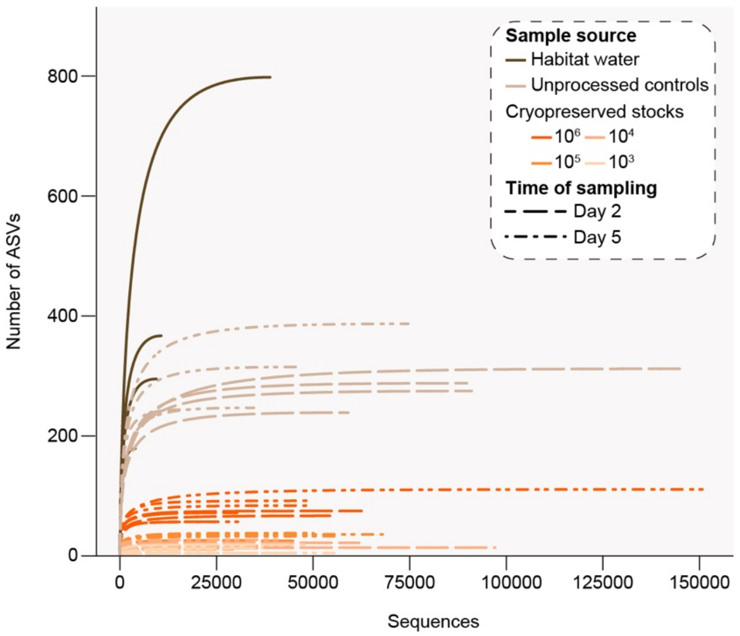


**Supplementary Fig. 5**. Rarefaction data from Illumina sequencing of water samples from a naturally occurring mosquito larval habitat in the field and resulting experimental microcosms. Reads from each water library were sampled starting at 1 sequence per step and increased in increments of 100 until the total number of reads per sample was reached. Lines are colored by sample source (rearing pans, brown; experimental microcosms containing unprocessed water, tan; experimental microcosms containing water plus material from a given cryopreserved stock, orange). Time of sampling of experimental microcosms is designated by line type (Day 2, long-dash; Day 5, dot-dash).

Supplementary Figure 6

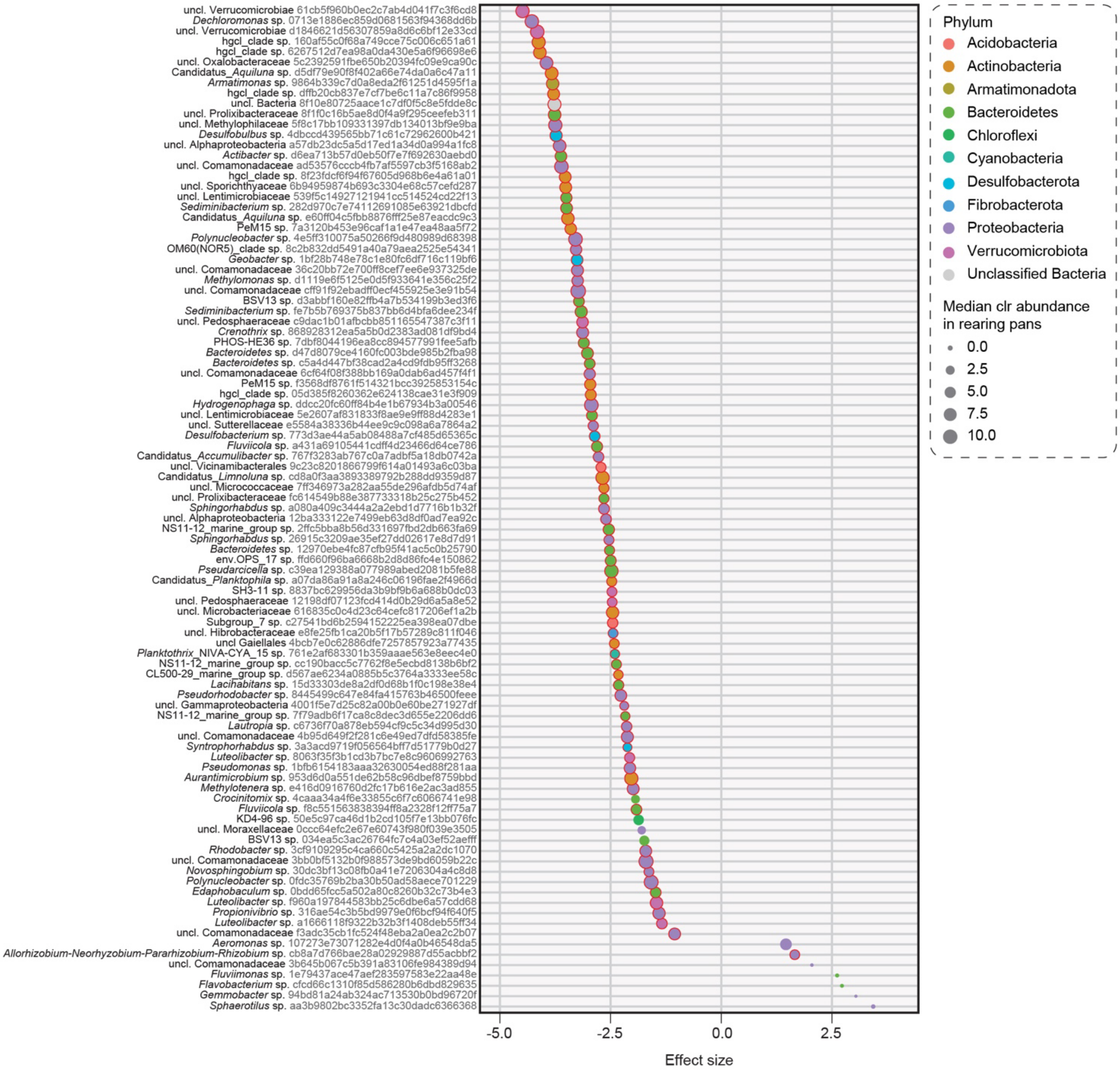


**Supplementary Fig. 6**. Differentially abundant ASVs in water samples collected from a naturally occurring mosquito larval habitat in the field and experimental microcosms containing unprocessed habitat water, identified through ALDEx2 testing (ALDEx2, $P < 0.05$; FDR, $P < 0.05$). Each ASV is presented with its lowest annotated taxonomic rank (to genus level) together with its ASV ID. The ASVs are color-coded according to the phyla they belong to and plotted according to their effect size, calculated as the levels in samples from experimental microcosms relative to levels in habitat water samples. Dot sizes correspond to the median clr abundance value for each ASV across habitat water samples. Dots with a bold red outline represent ASVs that were differentially abundant in both experimental microcosms containing unprocessed water and experimental microcosms containing water plus material from cryopreserved stocks (see Supplementary Fig. 7).

Supplementary Figure 7

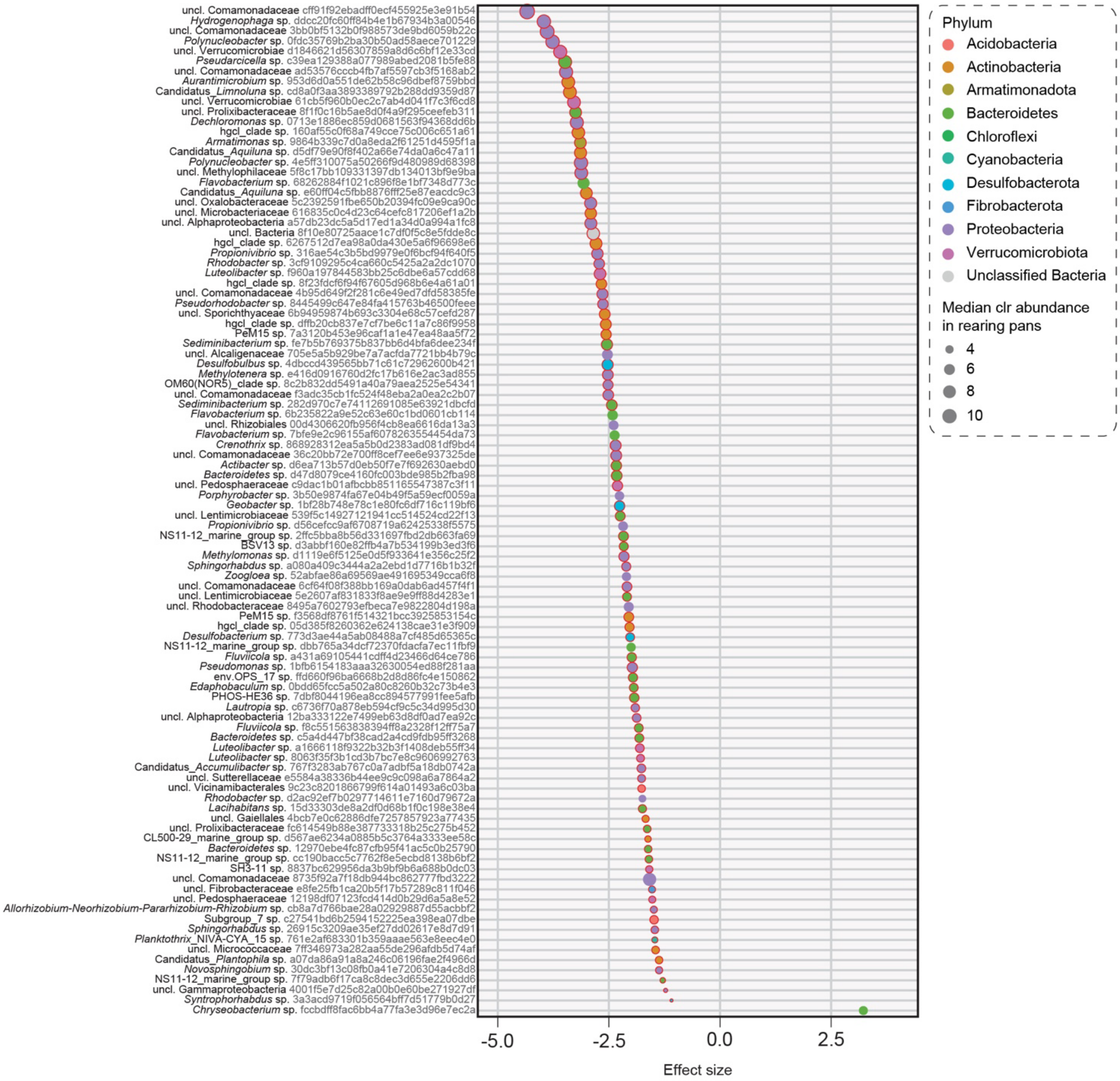


**Supplementary Fig. 7**. Differentially abundant ASVs in water samples collected from a naturally occurring mosquito larval habitat in the field and experimental microcosms containing water plus material from cryopreserved stocks, identified through ALDEx2 testing (ALDEx2, $P < 0.05$; FDR, $P < 0.05$). Each ASV is presented with its lowest annotated taxonomic rank (to genus level) together with its ASV ID. The ASVs are color-coded according to the phyla they belong to and plotted according to their effect size, calculated as the levels in samples from experimental microcosms relative to levels in habitat water samples. Dot sizes correspond to the median clr abundance value for each ASV across habitat water samples. Dots with a bold red outline represent ASVs that were differentially abundant in both experimental microcosms containing water plus material from cryopreserved stocks and experimental microcosms containing unprocessed water (see Supplementary Fig. 6).

### Supplementary Table 1

**Supplementary Table 1.** Sequencing and diversity statistics for 16S rRNA gene amplicon libraries prepared from water collected from conventional larval rearing pans in the laboratory and resulting experimental microcosms. Asterisks (*) indicate samples that were removed from the dataset prior to downstream analyses.

| Sample ID | Sample type | Density | Time of sampling | Total reads | Total ASVs | Shannon index |
|---|---|---|---|---|---|---|
| 50ml-A | Rearing pan | - | - | 43265 | 61 | 4.1742 |
| 50ml-B | Rearing pan | - | - | 27500 | 65 | 4.2197 |
| 50ml-C | Rearing pan | - | - | 71307 | 58 | 3.8723 |
| 50ml-D | Rearing pan | - | - | 45225 | 70 | 4.2343 |
| 50x-d2-A | Microcosm (unprocessed) | - | Day 2 | 122102 | 376 | 5.3964 |
| 50x-d2-B | Microcosm (unprocessed) | - | Day 2 | 60103 | 94 | 4.5934 |
| 50x-d2-C | Microcosm (unprocessed) | - | Day 2 | 112373 | 440 | 5.5159 |
| 50x-d2-D | Microcosm (unprocessed) | - | Day 2 | 52752 | 85 | 3.9012 |
| 50x-d5-A | Microcosm (unprocessed) | - | Day 5 | 8888 | 47 | 3.1674 |
| 50x-d5-B | Microcosm (unprocessed) | - | Day 5 | 25877 | 93 | 4.3186 |
| 50x-d5-C | Microcosm (unprocessed) | - | Day 5 | 16892 | 63 | 3.4301 |
| 50x-d5-D | Microcosm (unprocessed) | - | Day 5 | 7124 | 64 | 4.4214 |
| 0-d2-A | Microcosm (cryopreserved) | $10^8$ cells per ml | Day 2 | 998 | 17 | 2.8264 |
| 0-d2-B | Microcosm (cryopreserved) | $10^8$ cells per ml | Day 2 | 56898 | 50 | 4.0687 |
| 0-d2-C | Microcosm (cryopreserved) | $10^8$ cells per ml | Day 2 | 50352 | 53 | 3.8461 |
| 0-d2-D | Microcosm (cryopreserved) | $10^8$ cells per ml | Day 2 | 50979 | 57 | 4.2447 |
| 0-d5-A | Microcosm (cryopreserved) | $10^8$ cells per ml | Day 5 | 20128 | 45 | 3.5096 |
| 0-d5-B | Microcosm (cryopreserved) | $10^8$ cells per ml | Day 5 | 36407 | 57 | 3.9757 |
| 0-d5-C | Microcosm (cryopreserved) | $10^8$ cells per ml | Day 5 | 53569 | 48 | 3.6420 |
| 0-d5-D | Microcosm (cryopreserved) | $10^8$ cells per ml | Day 5 | 26264 | 61 | 4.2549 |
| 1-d2-A | Microcosm (cryopreserved) | $10^7$ cells per ml | Day 2 | 44633 | 38 | 3.3959 |
| 1-d2-B | Microcosm (cryopreserved) | $10^7$ cells per ml | Day 2 | 67859 | 46 | 3.5527 |
| 1-d2-C | Microcosm (cryopreserved) | $10^7$ cells per ml | Day 2 | 62482 | 45 | 3.5508 |
| 1-d2-D | Microcosm (cryopreserved) | $10^7$ cells per ml | Day 2 | 56016 | 40 | 3.6281 |
| 1-d5-A | Microcosm (cryopreserved) | $10^7$ cells per ml | Day 5 | 52662 | 47 | 3.6479 |
| 1-d5-B | Microcosm (cryopreserved) | $10^7$ cells per ml | Day 5 | 20921 | 49 | 3.7809 |
| 1-d5-C | Microcosm (cryopreserved) | $10^7$ cells per ml | Day 5 | 33806 | 44 | 3.5116 |
| 1-d5-D | Microcosm (cryopreserved) | $10^7$ cells per ml | Day 5 | 35559 | 52 | 3.7764 |
| 2-d2-A | Microcosm (cryopreserved) | $10^6$ cells per ml | Day 2 | 59358 | 37 | 3.1160 |
| 2-d2-B | Microcosm (cryopreserved) | $10^6$ cells per ml | Day 2 | 43085 | 42 | 3.6196 |
| 2-d2-C | Microcosm (cryopreserved) | $10^6$ cells per ml | Day 2 | 67607 | 41 | 3.3986 |
| 2-d2-D* | Microcosm (cryopreserved) | $10^6$ cells per ml | Day 2 | 28 | - | - |
| 2-d5-A | Microcosm (cryopreserved) | $10^6$ cells per ml | Day 5 | 61484 | 39 | 3.4294 |
| 2-d5-B | Microcosm (cryopreserved) | $10^6$ cells per ml | Day 5 | 37861 | 45 | 3.6205 |
| 2-d5-C | Microcosm (cryopreserved) | $10^6$ cells per ml | Day 5 | 36451 | 37 | 3.2768 |
| 2-d5-D | Microcosm (cryopreserved) | $10^6$ cells per ml | Day 5 | 27671 | 40 | 3.5898 |
| 3-d2-A | Microcosm (cryopreserved) | $10^5$ cells per ml | Day 2 | 31563 | 29 | 3.3386 |
| 3-d2-B | Microcosm (cryopreserved) | $10^5$ cells per ml | Day 2 | 53234 | 37 | 3.1552 |
| 3-d2-C | Microcosm (cryopreserved) | $10^5$ cells per ml | Day 2 | 38611 | 33 | 3.5208 |
| 3-d2-D | Microcosm (cryopreserved) | $10^5$ cells per ml | Day 2 | 61732 | 39 | 3.4932 |
| 3-d5-A | Microcosm (cryopreserved) | $10^5$ cells per ml | Day 5 | 57258 | 35 | 3.4729 |
| 3-d5-B | Microcosm (cryopreserved) | $10^5$ cells per ml | Day 5 | 41448 | 43 | 3.6010 |
| 3-d5-C | Microcosm (cryopreserved) | $10^5$ cells per ml | Day 5 | 34395 | 35 | 3.2676 |
| 3-d5-D | Microcosm (cryopreserved) | $10^5$ cells per ml | Day 5 | 32762 | 40 | 3.5156 |
| 4-d2-A | Microcosm (cryopreserved) | $10^4$ cells per ml | Day 2 | 51176 | 31 | 3.2377 |
| 4-d2-B | Microcosm (cryopreserved) | $10^4$ cells per ml | Day 2 | 49723 | 31 | 3.2008 |
| 4-d2-C* | Microcosm (cryopreserved) | $10^4$ cells per ml | Day 2 | 41 | - | - |
| 4-d2-D | Microcosm (cryopreserved) | $10^4$ cells per ml | Day 2 | 62944 | 37 | 3.4669 |
| 4-d5-A | Microcosm (cryopreserved) | $10^4$ cells per ml | Day 5 | 38359 | 32 | 3.4570 |
| 4-d5-B | Microcosm (cryopreserved) | $10^4$ cells per ml | Day 5 | 34889 | 31 | 3.3242 |
| 4-d5-C | Microcosm (cryopreserved) | $10^4$ cells per ml | Day 5 | 63315 | 34 | 3.3751 |
| 4-d5-D | Microcosm (cryopreserved) | $10^4$ cells per ml | Day 5 | 25329 | 37 | 3.4865 |
| 5-d2-A | Microcosm (cryopreserved) | $10^3$ cells per ml | Day 2 | 54637 | 27 | 3.1312 |
| 5-d2-B | Microcosm (cryopreserved) | $10^3$ cells per ml | Day 2 | 57438 | 30 | 3.0905 |
| 5-d2-C | Microcosm (cryopreserved) | $10^3$ cells per ml | Day 2 | 60946 | 29 | 3.2835 |
| 5-d2-D | Microcosm (cryopreserved) | $10^3$ cells per ml | Day 2 | 44747 | 27 | 3.3447 |
| 5-d5-A | Microcosm (cryopreserved) | $10^3$ cells per ml | Day 5 | 35667 | 30 | 3.2233 |
| 5-d5-B | Microcosm (cryopreserved) | $10^3$ cells per ml | Day 5 | 43450 | 37 | 3.1519 |
| 5-d5-C | Microcosm (cryopreserved) | $10^3$ cells per ml | Day 5 | 52643 | 32 | 3.2932 |
| 5-d5-D | Microcosm (cryopreserved) | $10^3$ cells per ml | Day 5 | 45081 | 32 | 3.4238 |

### Supplementary Table 2

**Supplementary Table 2.** Sequencing and diversity statistics for 16S rRNA gene amplicon libraries prepared from water collected from a naturally occurring mosquito larval habitat in the field and resulting experimental microcosms.

| Sample ID | Sample type | Density | Time of sampling | Total reads | Total ASVs | Shannon index |
|---|---|---|---|---|---|---|
| F-50ml-A | Habitat water | - | - | 3964 | 179 | 6.1297 |
| F-50ml-B | Habitat water | - | - | 38836 | 798 | 7.2439 |
| F-50ml-C | Habitat water | - | - | 9463 | 295 | 6.5045 |
| F-50ml-D | Habitat water | - | - | 10684 | 367 | 6.6975 |
| F-50x-d2-A | Microcosm (unprocessed) | - | Day 2 | 145719 | 312 | 5.6223 |
| F-50x-d2-B | Microcosm (unprocessed) | - | Day 2 | 90965 | 275 | 5.8426 |
| F-50x-d2-C | Microcosm (unprocessed) | - | Day 2 | 89703 | 288 | 5.8295 |
| F-50x-d2-D | Microcosm (unprocessed) | - | Day 2 | 59059 | 239 | 5.8823 |
| F-50x-d5-A | Microcosm (unprocessed) | - | Day 5 | 45717 | 315 | 6.0728 |
| F-50x-d5-B | Microcosm (unprocessed) | - | Day 5 | 17268 | 244 | 6.7062 |
| F-50x-d5-C | Microcosm (unprocessed) | - | Day 5 | 74564 | 387 | 5.9063 |
| F-50x-d5-D | Microcosm (unprocessed) | - | Day 5 | 34632 | 247 | 5.4338 |
| F-0-d2-A | Microcosm (cryopreserved) | $10^6$ cells per ml | Day 2 | 62480 | 75 | 3.5792 |
| F-0-d2-B | Microcosm (cryopreserved) | $10^6$ cells per ml | Day 2 | 54280 | 67 | 3.4459 |
| F-0-d2-C | Microcosm (cryopreserved) | $10^6$ cells per ml | Day 2 | 24488 | 57 | 3.2725 |
| F-0-d2-D | Microcosm (cryopreserved) | $10^6$ cells per ml | Day 2 | 30310 | 72 | 3.9956 |
| F-0-d5-A | Microcosm (cryopreserved) | $10^6$ cells per ml | Day 5 | 48528 | 92 | 3.6742 |
| F-0-d5-B | Microcosm (cryopreserved) | $10^6$ cells per ml | Day 5 | 30520 | 57 | 3.3095 |
| F-0-d5-C | Microcosm (cryopreserved) | $10^6$ cells per ml | Day 5 | 152484 | 111 | 3.5537 |
| F-0-d5-D | Microcosm (cryopreserved) | $10^6$ cells per ml | Day 5 | 48067 | 84 | 4.0770 |
| F-1-d2-A | Microcosm (cryopreserved) | $10^5$ cells per ml | Day 2 | 20678 | 21 | 3.0125 |
| F-1-d2-B | Microcosm (cryopreserved) | $10^5$ cells per ml | Day 2 | 46650 | 25 | 3.2540 |
| F-1-d2-C | Microcosm (cryopreserved) | $10^5$ cells per ml | Day 2 | 18065 | 25 | 2.9398 |
| F-1-d2-D | Microcosm (cryopreserved) | $10^5$ cells per ml | Day 2 | 34546 | 26 | 2.5901 |
| F-1-d5-A | Microcosm (cryopreserved) | $10^5$ cells per ml | Day 5 | 49969 | 38 | 2.9305 |
| F-1-d5-B | Microcosm (cryopreserved) | $10^5$ cells per ml | Day 5 | 57471 | 36 | 3.0436 |
| F-1-d5-C | Microcosm (cryopreserved) | $10^5$ cells per ml | Day 5 | 67926 | 36 | 2.7067 |
| F-1-d5-D | Microcosm (cryopreserved) | $10^5$ cells per ml | Day 5 | 57401 | 33 | 2.4124 |
| F-2-d2-A | Microcosm (cryopreserved) | $10^4$ cells per ml | Day 2 | 29184 | 11 | 1.7388 |
| F-2-d2-B | Microcosm (cryopreserved) | $10^4$ cells per ml | Day 2 | 61850 | 22 | 2.8832 |
| F-2-d2-C | Microcosm (cryopreserved) | $10^4$ cells per ml | Day 2 | 97008 | 14 | 1.4067 |
| F-2-d2-D | Microcosm (cryopreserved) | $10^4$ cells per ml | Day 2 | 36429 | 13 | 2.4710 |
| F-2-d5-A | Microcosm (cryopreserved) | $10^4$ cells per ml | Day 5 | 44047 | 16 | 2.2003 |
| F-2-d5-B | Microcosm (cryopreserved) | $10^4$ cells per ml | Day 5 | 41637 | 27 | 2.4174 |
| F-2-d5-C | Microcosm (cryopreserved) | $10^4$ cells per ml | Day 5 | 35402 | 13 | 1.8279 |
| F-2-d5-D | Microcosm (cryopreserved) | $10^4$ cells per ml | Day 5 | 35949 | 17 | 2.7682 |
| F-3-d2-A | Microcosm (cryopreserved) | $10^3$ cells per ml | Day 2 | 43035 | 5 | 0.0089 |
| F-3-d2-B | Microcosm (cryopreserved) | $10^3$ cells per ml | Day 2 | 36831 | 13 | 0.7838 |
| F-3-d2-C | Microcosm (cryopreserved) | $10^3$ cells per ml | Day 2 | 47713 | 19 | 1.8455 |
| F-3-d2-D | Microcosm (cryopreserved) | $10^3$ cells per ml | Day 2 | 15000 | 5 | 0.0602 |
| F-3-d5-A | Microcosm (cryopreserved) | $10^3$ cells per ml | Day 5 | 56109 | 5 | 0.0455 |
| F-3-d5-B | Microcosm (cryopreserved) | $10^3$ cells per ml | Day 5 | 24035 | 9 | 1.9925 |
| F-3-d5-C | Microcosm (cryopreserved) | $10^3$ cells per ml | Day 5 | 34929 | 19 | 1.4651 |
| F-3-d5-D | Microcosm (cryopreserved) | $10^3$ cells per ml | Day 5 | 47450 | 5 | 1.4273 |
